# Early Amyloid-β Toxicity Disrupts Nervous System Connectivity During Aging in *Caenorhabditis elegans*

**DOI:** 10.64898/2026.07.27.740981

**Authors:** Dilip Kumar Yadav, Christopher W Connor, Christopher V Gabel

## Abstract

Alzheimer’s disease (AD) is characterized by progressive functional neuronal decline ultimately resulting in severe cognitive impairment. However, how neuron function is altered at the cellular level during the critical early stages of the disease and how disease progression relates to the process of normal neuronal aging are poorly understood. To address these fundamental questions, we performed comprehensive multi-neuron imaging in *Caenorhabditis elegans* (*C. elegans*) with pan-neuronal expression of human amyloid β_1-42_ peptide (nAβ), the major plaque forming peptide in early AD progression. Measuring neuron activity, connectivity and system wide dynamics with single cell resolution across the *C. elegans* lifespan, we compare Aβ-associated neuronal dysfunction with that of normal aging. Our experiments revealed that nAβ expressing animals exhibit a unique loss of positively correlated neuron connectivity, premature disruption of system wide dynamics, and reduced overall neuronal activity. These neuronal effects correspond to premature impairments in behavior including mechanosensory response as well as the animal’s ability to navigate its environment (*i.e*., chemotaxis and thermotaxis). The effects of nAβ expression are distinct from, and in addition to, the process of normal aging that is characterized by a progressive loss of anti-correlated (*i.e.* inhibitory) neuronal signaling and slower behavioral decline. An increase in resistance to aldicarb, an acetylcholinesterase inhibitor, as well as downregulation of key one-carbon metabolism (OCM) genes (*metr-1, sams-1*, involved in choline metabolism, the precursor of acetylcholine synthesis) implicate compromised acetylcholine-mediated excitatory transmission in nAβ expressing worms. Supplementation with OCM metabolites (choline, methionine, cysteine) in nAβ expressing worms improved behavior and helped restore OCM gene expression as well as levels of Ach signaling. Likewise, perturbation of the serine synthesis pathway (SSP) that links glycolysis to OCM, altered Ach signaling and OCM gene expression in nAβ animals. Genetic mutations that directly regulate excitatory/inhibitory balance neuronal signaling (*unc-2*/CaV2α) also help to reduce nAβ-associated behavioral deficits, while NMDA receptor (*nmr-1*) mutants showed neither protective effect nor it directly rescue the loss of positive neuron correlativity unique to nAβ animals. Our results demonstrate the unique effects of nAβ toxicity on neuronal dynamics and connectivity as well as the role of key metabolic pathways within the context of a complete, intact, aging nervous system.

## Introduction

In its later stages, Alzheimer’s Disease (AD) results in severe cognitive impairment and neuronal death that effects millions of people particularly within the aging population. However, the process of neuronal dysfunction and decline starts well before obvious clinical symptoms (1). Early AD pathologies at the cellular level include instability of neuronal firing rates and synaptic signaling (2,3). Elucidating these early steps of functional neuronal decline at the cellular level is fundamental to understanding the underlying cellular pathology and ultimately formulating effective therapeutic strategies. However, the picture is further complicated by aging, the primary risk factor for AD, that also incurs neuronal decline altering activity, synaptic connectivity and system dynamics (4). Disentangling the early decline of the disease state from that if normal aging and understanding the interdependencies of these two processes remains a critical challenge to ultimately mitigating the cellular progression of AD.

Substantial evidence links amyloid-β peptide (Aβ) toxicity to neuronal dysfunction well before evidence of clear clinical symptoms (5), including studies measuring the breakdown of brain network signaling (6). At the cellular level the early stages of AD have been linked to increased plaque formation of Aβ aggregates, neuronal hyperexcitability, as well as degradation of both cholinergic and glutaminergic synaptic connections (7). Current hypotheses suggest that it is a breakdown of core homeostatic machinery leading to dysregulation of neuronal activity that drive AD pathogenesis (3). However, the cellular mechanisms of this degenerative process and its relation to normal aging have yet to be fully understood. It is therefore critical to uncouple and understand the toxic effects of Aβ on neuronal function at the cellular and circuit level within the context of normal aging.

Recent human imaging studies reveal changes in brain circuit dynamics that correspond to functional decline of cognitive tasks in AD (8). However, such large-scale brain measurements (such as fMRI, and EEG) that are possible in humans and other mammals lack the resolution necessary to measure functional changes at the cellular level. By contrast, the simple nematode worm *Caenorhabditis elegans* (*C. elegans*) offers a powerful model to investigate functional neuronal decline on the cellular level due to its simple, completely mapped nervous system, short lifespan, and genetic tractability (9). Recent advances in multi-neuron fluorescence microscopy have enabled real-time measurement of neuronal activity across the majority of the *C. elegans* nervous system with single cell resolution. This unprecedented scope provides insights into neuronal function, and connectivity within the context of system-wide dynamics of an intact nervous system (10). Employing these techniques, we have recently imaged the functional decline of the *C. elegans* nervous system during normal, healthy aging (Wirak et al, 2022), revealing conserved mechanisms of neuronal aging, including increased hyperactivity that mimic changes in neuronal excitability measured in the mammalian brain, and a loss of inhibitory signaling also observed in mammals in various context (11,12). Moreover, many genes underlying cellular mechanisms of neuronal aging and AD pathology are highly conserved in *C. elegan*s. Numerous studies of Aβ toxicity have revealed conserved mechanisms of stress response across human, mouse and *C. elegans* (9). Transgenic *C. elegans* with pan-neuronal expression of low-levels of human Aβ_1-42_ peptide display accelerated behavioral decline as well as neurodegeneration with age that is dependent on mitochondrial breakdown and oxidative stress paralleling findings in mammals (13). Thus *C. elegans* recapitulates many of the central mechanisms of neuronal degradation in Aβ models of AD and normal aging.

In this study, we employ the competence and simplicity of *C. elegans* to elucidate the effects of Aβ neurotoxicity on nervous system dynamics within the context of aging. Comprehensive neuronal imaging in wild-type and nAβ expressing animals across the animals life-span allows us to isolate and understand changes in neuron activity, connectivity and system-wide dynamics within the normal and pathological states. Exploiting the genetic tractability of *C. elegans*, we test the relevance and potential influence of metabolic pathways in these processes of neuronal degradation. Our results demonstrate how Aβ toxicity alters nervous system function at the cellular and circuit level and how this differs from the effects of normal aging, thus helping to reveal the critical first steps of neuronal decline in AD.

## Results

### *C. elegans* neurons with Human Amyloid Beta (Aβ_1-42_) expression exhibit unique decline of nervous-system dynamics

In this study we employed an established *C. elegans* model of AD consisting of pan-neuronal expression of human amyloid beta peptide (Aβ_1-42_) under a temperature sensitive promoter (strain CL2355) that shows progressive neurodegeneration (14). To enable multi-neuronal imaging the Aβ_1-42_ strain was crossed with a neuronal imaging strain (QW1217;OH15263), with pan-neuronal expression of nuclear-localized tagRFP and cytoplasmic GCaMP6s (See Methods, Wirak et al 2022) (11). Temperature dependent amyloid beta peptide expression allowed us to rear the animals through the initial stages of development with minimal or no pan-neuronal Aβ (nAβ) expression at 16°C. By transitioning animals to higher temperature (23°C), at early young adults/day1 age (illustrated in Supp Figure 1a), we induced nAβ expression throughout adulthood.

**Fig. 1.**
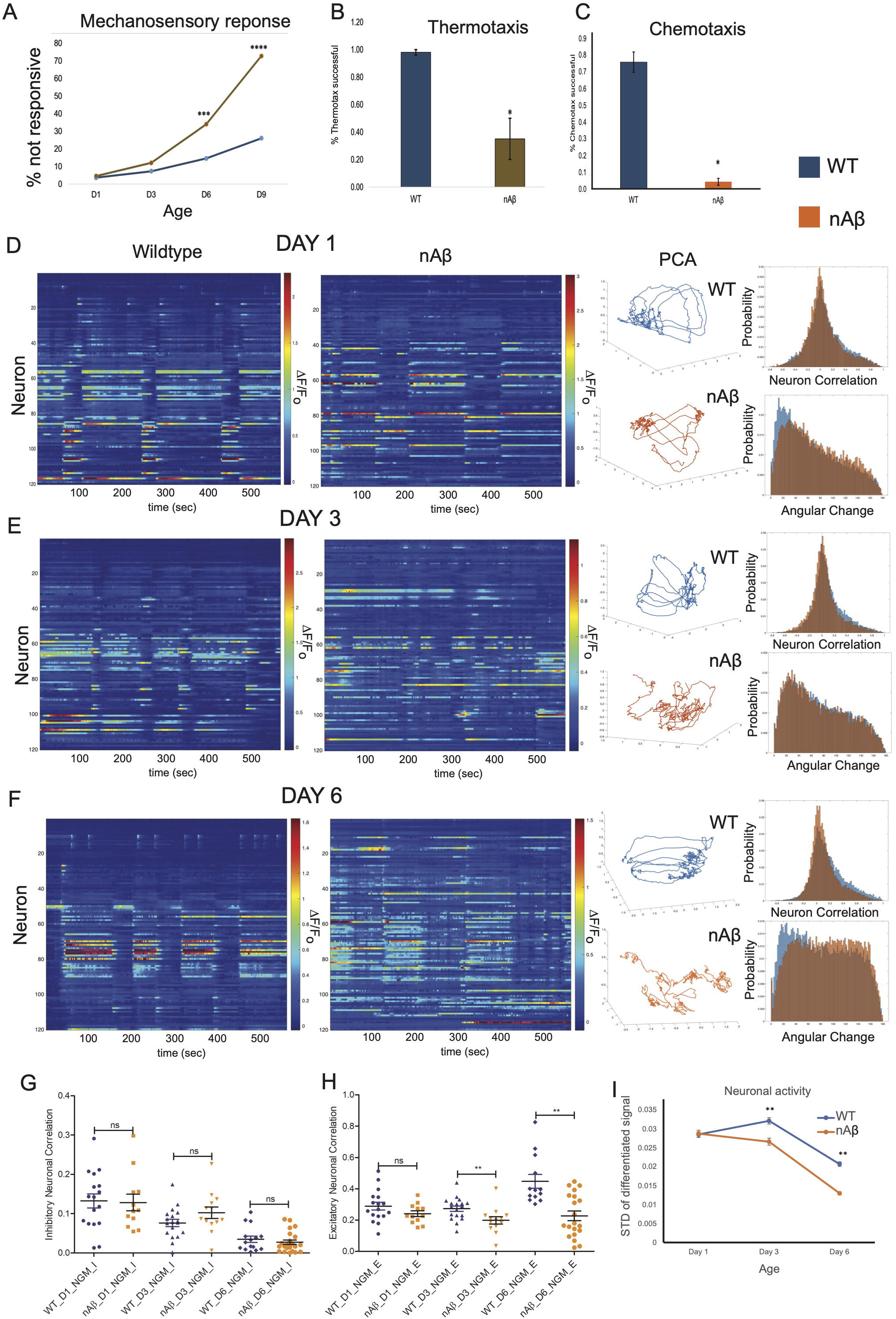
The loss of excitatory signaling in nAβ-mediated neuronal dysfunction. (A). Anterior touch response plotted as percentage of animals not responding (NR). Wildtype (WT) (QW1217;OH15263) and pan neuronal Abeta (nAβ) expressing (CL2355; QW1217;OH15263) animals were tested across different ages (day1, day3, day6 and day9). (B). Thermotaxis and (C). Chemotaxis behavior assay in WT and nAβ worms on day6. (D). Day1, (E). Day 3, and (F). Day 6, Left panel shown example heatmaps of 120-neuron GCaMP fluorescence measurements, captured in the head region of WT and nAβ worm. Middle panels show, Three-dimensional plots displaying the trajectories of the first three principal components derived from the displayed example trials (WT in blue, nAβ in orange). Right panels, aggregate probability histograms of neuron pair correlativity among the 40 most dynamically active neurons, in WT (blue) and nAβ worms (orange) and aggregate probability histograms of the angular directional changes in principal component trajectories for all animals measured at various ages (day1, day3 and day6) in WT (blue) and nAβ worms (orange). Comparison of inhibitory (G), and excitatory (H) neuronal correlation between WT (blue) and nAβ (orange) worms on day1, day3 and day6. (I) Overall neuronal activity between control and nAβ worms measured as mean of standard deviation of spontaneous neuronal activity on day1, day3 and day6. All animals were initially cultured at 16°C and then switch to 23°C from day 1 onwards to induce nAβ expression in adults only (see methods). pValue * < 0.05, **<0.01, ***<0.001.

*C. elegans* displayed progressive decline of sensory response behaviors with age. We quantified this by measuring mechanosensory response to a light touch stimulus (eyelash touch) and found that nAβ expressing animals show a premature decline in touch response compared to age matched wild type controls (Fig. 1A). Notably, we did not measure changes in mechanosensory response in nAβ worms exposed to an intermediate temperature (20°C) until later in adulthood, day 9 (Supp fig. 1b). We also measured the animals’ ability to seek out preferable temperature (thermotaxis) and chemical signals (chemotaxis) while crawling on an agar plate. We find that nAβ expressing animals are significantly impaired in their ability to perform these behaviors compared to controls at adult day 6 (Fig. 1B, and 1C). These experiments established a significant behavioral defect in animals reared as adults at 23°C with pan-neuron Aβ_1-42_expression.

To investigate the effects of Aβ_1-42_ expression on neuronal function at the cellular, circuit and system levels, we performed comprehensive multi-neuron imaging in both WT and nAβ expressing animals. Animals were imaged at progressive ages, adult day 1, day 3 and day 6 for 10 min trials of spontaneous activity. Wild-type *C. elegans* typically live for ∼2 weeks but physical degradation of neurons in nAβ expressing animals prevented effective neuronal imaging past adult day 6. As described in the methods we employed fluorescence light sheet microscopy to capture calcium sensitive GCaMP6s signal from 120 neurons within the *C. elegans* head ganglion region. This technique allows for optical measurement of neuron activity (via calcium dynamics) across a majority of neurons in the animal’s head with single cell resolution (11).

Example imaging trials of wildtype (QW1217;OH15263) and pan-neuronal Aβ expressing strain (nAβ,CL2355; QW1217;OH15263) worms are displayed as neuronal activity heatmaps for adult days 1,3,6 in Figure 1 (Fig. 1D-1F, left). Visualization of system-wide dynamics of the multi-neuron data sets through Principal Component Analysis (PCA) illustrated that, neuronal activity in healthy worms typically trace out a smooth looping trajectory in the PCA plots. This demonstrates organized and well-defined system-wide dynamics as the animals transition between distinct behavioral states (i.e. forward, backward crawling, Fig. 1D-1F, center). Studies have shown that such PCA trajectories correspond to the system occupying and smoothly transitioning between well-defined behavioral states (15). By contrast, PCA analysis of neuronal imaging in nAβ expressing worms display irregular and randomized trajectories in the PCA plots. To quantify this breakdown in system-wide dynamics across animals, we measured temporal continuity of the system as the smoothness of the PCA trajectories over time (as in Wirak et al 2022). A smoothly curving PCA trajectory is characterized by small angular changes in direction between successive time frames, while a completely stochastic, randomized trajectory will have equally distributed angular changes in all possible directions (over 180°). Plotting angular changes from all-time windows in all trials at one condition on a cumulative histogram, we see that day 1 WT animals are skewed toward smaller angular changes (Figure 1D, right bottom panel, blue), highlighting the smooth characteristic of the trajectories and temporal continuity of the system. With age, this bias is progressively lost, generating histograms with more equal distributions across all angular changes reflecting erratic randomly changing trajectories (Figure 1E and 1F, right bottom panel, blue). By this metric, nAβ expressing worms exhibit increasing stochastic, randomized trajectories as early as adult day 1 and are completely randomized by adult day 6 (Figure 1D-1F, right bottom panel, orange). The increased stochasticity of PCA traces reflects an accelerated loss of organization and temporal continuity of neuronal dynamics in nAβ expressing animals.

Performing quantitative analysis of cell-to-cell correlation of activity within the multi-neuron data sets, we can measure the degree of functional excitatory signaling, *i.e.* positively correlated neuron pairs, and inhibitory signaling, *i.e.* negatively correlated neuron pairs, within the system. Data is shown as probability histograms of correlativity between all neuron pairs imaged across all trials at that condition. They can be quantified by the percentage of neuron pairs that are highly correlated or anti-correlated (see methods, Figure 1D-F, right top panels, shows data by age for WT and nAβ). As in our previous study (Wirak et al 2022), WT animals display a progressive loss of negatively correlated neuron pairs with age while preserving positively correlated neuron pairs (Figure 1G-H, blue, Table 1) (11). We find that nAβ expressing animals exhibit a loss of negative correlativity similar to that of normal aging. However, they also display a relative loss of positively correlated neuron pairs (Figure 1G-H, orange, Table 1). nAβ animals raised at 20°C, with minimal Aβ expression, did not display any difference from WT under the same conditions. (Supp fig. 1c-d, Table 1). Thus, the loss of excitatory signaling appears to be an element of functional neuronal decline that is unique to nAβ expressing animals.

We measured the overall neuronal activity of the system as the mean activity across all neurons imaged at each condition. Neuron activity was measured as the standard deviation (STD) of the differentiated signal for each neuron’s fluorescence trace. Active neurons with widely dynamic traces will yield large STD and vice versa. As shown in Figure 1I, cumulative neuronal activity was significantly decreased in nAβ worms compared to WT in older animals (adult day 3, day 6). This reduction in overall system activity corresponds with the reduction in positively correlated, excitatory, neuron pairs observed in these animals with age.

Taken as a whole, these initial experiments demonstrate a unique profile of systemwide functional degradation in nAβ expressing animals, that is comprised of premature disruption of system state dynamics and organization, loss of excitatory signaling and a system-wide decrease in activity dynamics. These neuronal effects parallel the progressive decline of mechanosensory behavior, as well as more complex chemo- and thermo-taxis behaviors in these animals. Given these results, we focused additional experiments below on adult day 6 as the time point at which we could observe the most robust changes both in behavior and neuronal dynamics.

### Disruption of positive neuronal connectivity in human Amyloid beta (Aβ) expressing *C. elegans* is linked to One Carbon Metabolism (OCM) pathways

The loss of positive neuronal correlativity in nAβ expressing worms suggests a possible disruption in acetyl cholinergic (ACh) signaling, which is one of the primary excitatory neurotransmitters in the *C. elegans* nervous system (16). ACh signaling *via* synaptic transmission can be directly assayed in *C. elegans* by measuring the time to paralysis when the animals are placed on Aldicarb, an acetylcholinesterase inhibitor (17). We find that nAβ expressing worms take significantly longer time to paralyze compared to control worms when subjected to 1 mM Aldicarb, in accordance with a decrease of ACh signaling (Fig. 2A).

**Fig. 2.**
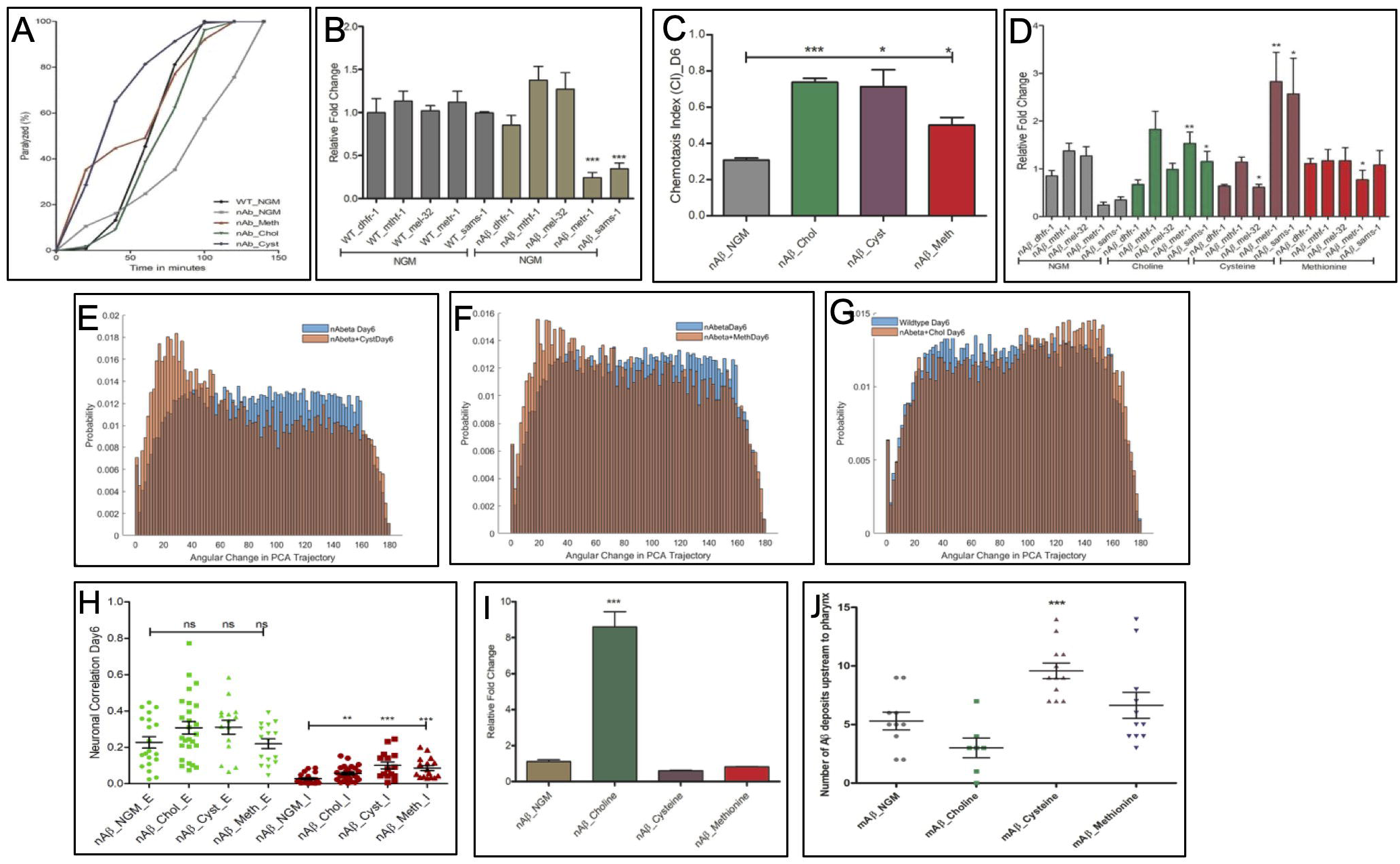
Role of OCM in Amyloid beta (Aβ) mediated neuronal decline. (A). Aldicarb assay of WT and nAβ worms (along with OCM metabolite supplementation in nAβ worms) on adult day6. (B). Quantitative RT-PCR of selected genes critical for OCM pathway in WT and nAβ worms (C). Chemotaxis behavior assay in nAβ worms on day6 with different OCM metabolite supplementation. (D) Effect of Choline, Cysteine and Methionine supplementation on the expression level of selected genes critical for OCM pathway in nAβ worms. Aggregate probability histograms of the angular directional changes in principal component trajectories for all nAβ worms on day6 measured upon various OCM metabolite supplementation (E) Cysteine, (F) Methionine and (G) Choline. (H). Quantitative comparison of excitatory (green) and inhibitory (red) signaling in nAβ worms on, upon various OCM metabolite supplementation. (I). Effect of Choline, Cysteine and Methionine on amyloid beta expression level in nAβ worms. (J). Effect of Choline, Cysteine and Methionine on amyloid beta plaques deposition in pharyngeal regions of worms expressing human amyloid beta in muscle cells of *C. elegans* (CL2006). All animals were adult day 6, pValue * < 0.05, **<0.01, ***<0.001.

Choline, a precursor of acetylcholine synthesis, is directly linked to One Carbon Metabolism (OCM) (18), a central metabolic pathway that has been clinically linked to age associated neurodegenerative diseases including Alzheimer’s disease in humans (19–21). Using quantitative RT-PCR, we measured that two critical OCM genes, Methionine Synthase (*metr-1*) and S-Adenosyl Methionine Synthetase (*sams-1*), are significantly down regulated in nAβ worms compared to control animals (Fig. 2B). Likewise, S-Adenosyl Methionine Synthetase mutant worms (*sams-1*) exhibit delayed paralysis in the aldicarb assay, indicating decreased ACh signaling in genetically OCM perturbed worms (Supp Fig. 2a). To further test the role of ACh signaling and OCM metabolism in Aβ associated decline, we exposed animals to OCM supplements Choline, Methionine and Cysteine (from day1-day6). We found that these supplements significantly improve the chemotaxis behavior of nAβ worms (on day 6, Fig. 2C) and did not affect WT (Supp Fig 2b). Choline also improves chemotaxis ability in the *sams-1* background (Supp Fig 2c). OCM supplements also rescued the time to paralysis on Aldicarb in nAβ expressing animals, indicating increased ACh signaling (Fig. 2A) and restore the expression level of OCM genes *metr-1* and *sams-1* in nAβ worms (Fig. 2D). There was no effect of these supplements on *metr-1* and *sams-1* expression in the WT animals, although, we observed someother genes expression was altered with these supplementations (Supp Fig 2d).

The transulfuration pathway, which is downstream of OCM, also plays a critical role in neurodegenerative diseases by controlling oxidative stress response in degenerating neurons (22,23). We found that genes involved in this pathway were also differentially regulated in nAβ expressing worms and that most were restored with OCM supplementation in nAβ worms (Supp fig 2e-h). Taken as a whole, we find that nAβ expression modulates numerous OCM and related genes and conversely that OCM nutritional supplements incur restorative effects in nAβ expressing animals.

Performing multi-neuronal imaging in nAβ expressing animals with OCM supplementation, we found that elements of functional decline were altered by OCM metabolites. PCA analysis revealed that Cysteine supplementation had a positive effect on system dynamics helping to preserve temporal continuity (smoothness of the PCA trajectories, Fig.2E). However other supplements had minimal or no effect in this regard (Fig. 2F and 2G). Importantly, OCM supplements in nAβ expressing animals consistently helped to preserve neuron anti-correlativity that declines during normal aging. However, we did not measure any significant effect of these supplements on the loss of positive neuron correlativity that is unique to nAβ expressing animals (Fig. 2H, Table2). Supplementation of these metabolites in control WT worms had no effect on neuron correlativity at adult day 6 (Supp fig. 2i, Table2), with the exception of cysteine that displayed reduced positive correlativity. In addition, we tested Vitamin B12, another micronutrient which is critical for OCM and clinically associated with neurodegenerative diseases (24,25) but did not detect any effect on neuronal activity in nAβ worms (Supp fig. 2j, Table2).

Further, we tested the effect of OCM supplements on nAβ expression levels in our transgenic animals. qRT-PCR outcome shows minimal effects on Aβ levels with all supplements with the exception of choline that showed enhanced Aβ expression (Fig. 2I, WT animals showed no Aβ expression as expected, Supp fig 2k). Likewise, Aβ plaque deposition in transgenic animals (CL2006) expressing the Aβ_1-42_ construct in muscle cells (where plaques can be easily imaged and quantified via a thioflavin S fluorescent staining) was not largely altered by OCM supplements with the exception of cysteine that showed increased number of plaques (Fig. 2J, Table2 and Supp fig. 2l). Thus, the effects of OCM supplements do not appear to be caused by a lack of Aβ expression.

### Perturbation of the Serine Synthesis Pathway causes a loss of cellular signaling similar to nAβ expression

Studies in mammals have reported significant contradictory effects of the Serine Synthesis Pathway (SSP) on Alzheimer’s disease onset and progression at the molecular and cellular levels (26,27). As the SSP is directly upstream of OCM and links glycolysis to OCM, we investigated it in our *C. elegans* nAβ model. We found that blocking SSP (NCT502 treatment) in nAβ animals increased Aldicarb resistance consistent with a further loss of ACh signaling, while L-serine supplementation had the opposite effect returning Aldicarb sensitivity close to WT levels (Fig. 3A). Interestingly, in control WT animals blocking SSP with a pharmacological agent (NCT502), or supplementing it *via* L-serine, results in prolonged Aldicarb resistance suggesting reduced ACh signaling (Fig. 3B). We did not observe any effect on chemotaxis behavior either in control or nAβ worms under these conditions (Fig. 3C and 3D).

**Fig. 3.**
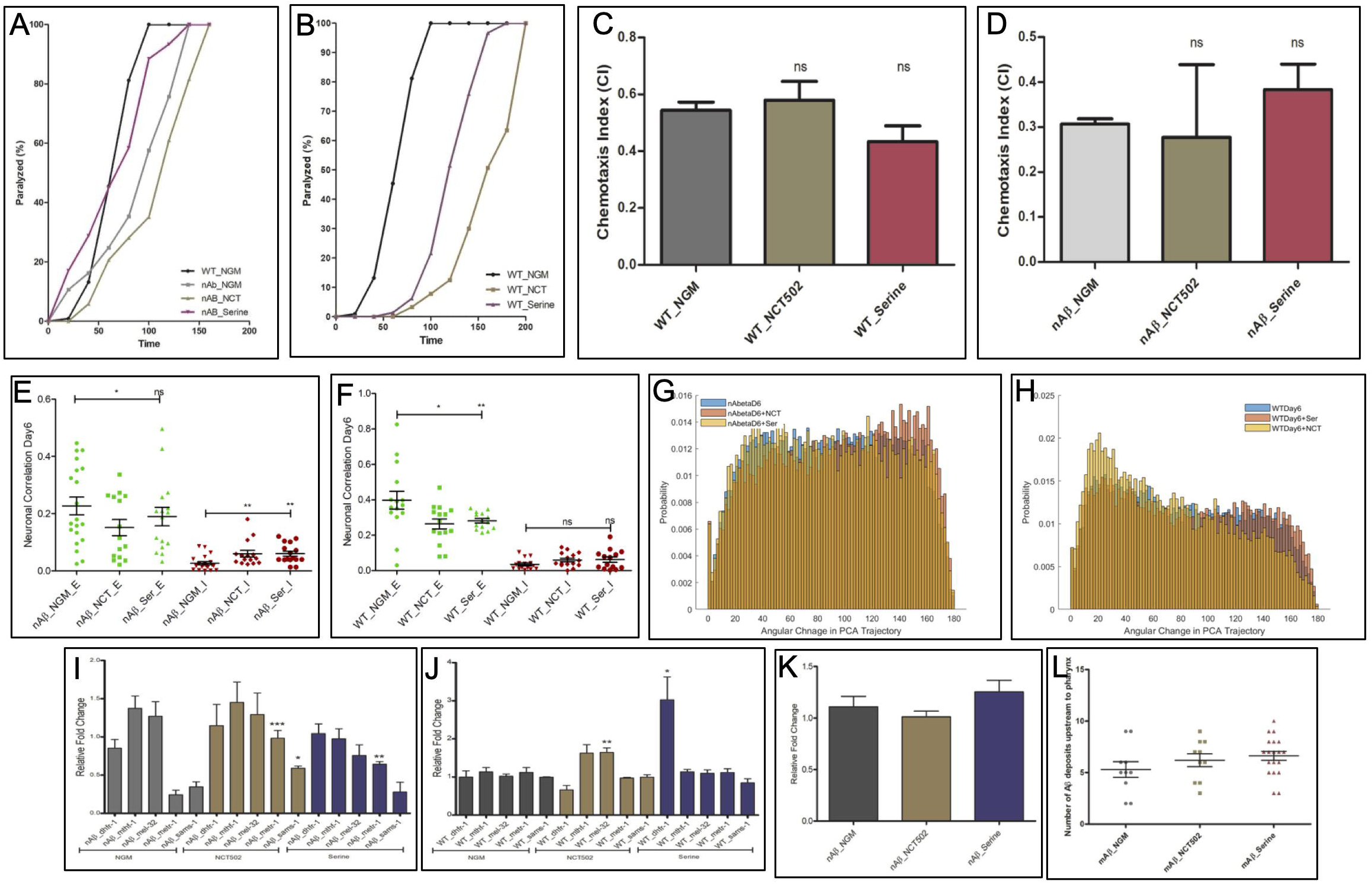
Effects of serine synthesis pathway modulation on nAβ-mediated neuronal dysfunction. The effect of perturbation of the serine synthesis pathway on acetyl choline signaling assessed by aldicarb assay in (A) nAβ worms and (B) WT worms. The effect of perturbation of the serine synthesis pathway on chemotaxis behavior in (C) WT and (D) nAβ worms. Quantitative comparison of excitatory (green) and inhibitory (red) signaling upon NCT502 and Serine supplementation in nAβ (E) and WT (F) worms. Aggregate probability histograms of the angular directional changes in principal component trajectories for all animals measured upon NCT502 and Serine supplementation in (G) nAβ and (H) WT worms. Effect of NCT502 and Serine supplementation on expression level of selected genes critical for OCM pathway in (I) nAβ and (J) WT worms. (K). Effect of NCT502 and Serine supplementation on amyloid beta expression level in nAβ worms. (R). Effect of NCT502 and Serine on amyloid beta plaque deposition in the muscle cells of the pharyngeal (head) regions of worms expressing human amyloid beta in muscle cells (CL2006). All animals were raised a 23 °C from day 1 of adulthood and tested at adult day 6. pValue * < 0.05, **<0.01, ***<0.001.

Multi-neuronal imaging revealed that manipulating the SSP either by blocking with NCT502, or supplementing with L-serine, helped to preserve the number of anticorrelated neuron pairs in nAβ animals at adult day 6 (Fig. 3E). However, we also found an increased loss of positive neuron correlation in most cases for both WT and nAβ animals (Fig. 3E, 3F, Table3) and no effect on temporal continuity of activity traces (with the exception of some improvement with NCT502 in WT) (Fig 3G,H). Molecular analysis revealed that NCT502 treatment helps to restore the expression levels of OCM genes *metr-1* and *sams-1* in nAβ worms although L-serine supplementation restored only *metr-1* (Fig. 3I). Whie *metr-1* and *sams-1* levels were not affected in control worms (Fig. 3J). We did not find any consistent significant expression differences of genes critical for SSP (*phgdh-1, psat-1* and *psph-1*) between control and nAβ worms (Supp fig. 3a), or with NCT502 or L-serine supplementation (with the exception of *psph-1* which showed increased expression upon NCT502 treatment in nAβ animals, Supp fig. 3b,c). Investigating the effect of SSP perturbation on the downstream transulfuration pathway, we find that numerous genes were differentially regulated in nAβ, and expression levels of most altered genes were further restored by NCT502 but not by L-serine treatments (Supp fig 3d-3f). There was no effect of NCT502 and L-serine either on the expression level of Aβ measured by qRT-PCR (Fig. 3K), nor in the number of Aβ plaques observed in muscle cells of worms (Fig. 3L, Table3 and Supp fig 3g-3h)

Serine racemase (*serr-1*), which is involved in conversion of L-serine to D-serine, is reported to have neurotoxic effects in Alzheimer’s disease, primarily by modulating N-methyl-D-aspartate (NMDA) receptors (28). We found that, while *serr-1* expression is not altered by nAβ expression alone, the SSP blocking agent NCT502 significantly increases the expression of *serr-1* in nAβ worms compared to controls, while L-serine supplementation significantly decreased *serr-1* in nAβ worms compared to controls (Supp fig 3i). Of the other OCM metabolites tested only Choline significantly increases the *serr-1* expression in nAβ worms compared to control (Supp fig 3j).

### Potential molecular mechanisms altering neuronal signaling in nAβ animals

To further explore the mechanism of loss of neuronal connectivity in aging nAβ worms, we employed the gain-of-function (gof) mutation in *unc-2*//CaV2α, (unc-2(zf35)). *unc-2* encodes the ortholog of the pore-forming alpha-1A subunit of the voltage-dependent P/Q-type calcium channel (CACNA1A) in *C. elegans*, and the gain-of-function mutation is known to potentiate excitatory/ACh signaling in the *C. elegans* nervous system (29). We found that while *unc-2(gof)* (QW1348) did not alter chemotaxis behavior at adult day 6 in a WT background, the *unc-2(gof)* mutation effectively recovered the chemotaxis defect of nAβ animals restoring it to WT levels (Fig. 4A). Performing multi-neuron imaging in nAβ animals, we found that the *unc-2(gof)* mutation slightly improved anti-correlated signaling but had no measurable effect on the degree of positive neuron correlativity (Fig. 4B, Table 4). In WT background *unc-2(gof)* worms show similar effects on day 6, with increased neuron anti-correlativity but no effect on positive correlativity compared to controls (Supp fig. 4a, Table 4).

**Fig. 4.**
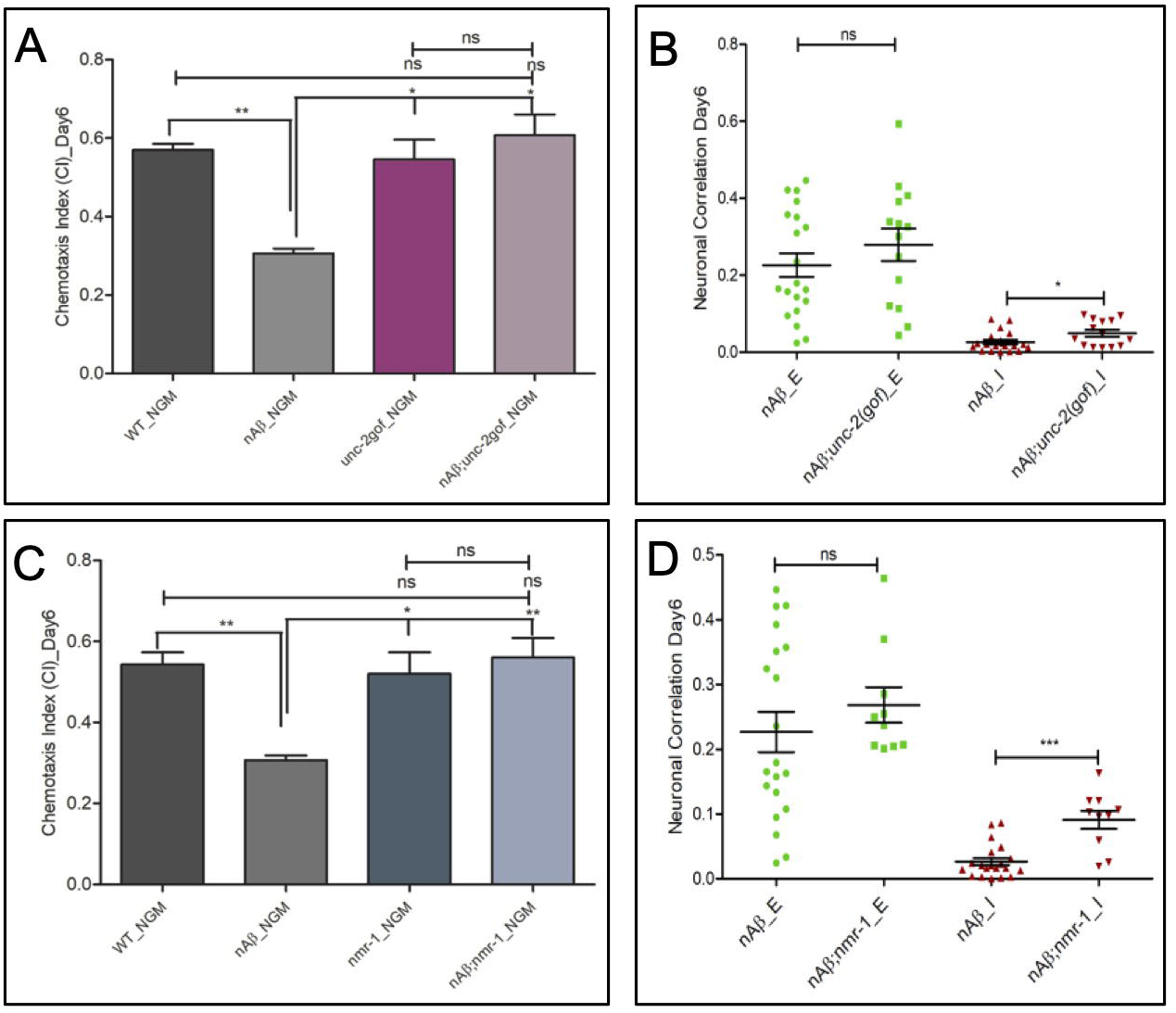
The molecular mechanism of Aβ-mediated excitatory signaling decline. (A). Chemotaxis behavior assay in control, nAβ, unc-2(gof) and nAβ;unc-2(gof) worms. (B). Quantitative comparison of excitatory (green) and inhibitory (red) neuronal signaling in nAβ and nAβ;unc-2(gof)) worms. (C). Chemotaxis behavior assay in WT, nAβ, *nmr-1* and nAβ*;nmr-1* worms. (D). Quantitative comparison of excitatory (green) and inhibitory (red) neuronal signaling in nAβ and nAβ;*nmr-1* worms. All animals were raised a 23 °C from day 1 of adulthood and tested at adult day 6. pValue * < 0.05, **<0.01, ***<0.001.

The NMDA (N-methyl-D-aspartate) glutamate receptor is an important excitatory signaling component that is thought to play a role in Alzheimer’s disease (AD) (30). Under healthy conditions, it supports synaptic plasticity, learning, and memory by mediating calcium influx (31,32). However, in AD, Aβ neurotoxicity is thought to cause excessive activation of NMDA leading to excitotoxicity, calcium overload, mitochondrial dysfunction, and oxidative stress, contributing to neuronal death (30–32). The NMDA receptor is well conserved in *C. elegans* (encoded by *nmr-1*) and plays a critical role in excitatory signaling within the animal’s command motor circuitry controlling behavioral states. Therefore, we investigated the role of the NMDA receptor in the nAβ-associated changes in neuronal dynamics. We found that *nmr-1* null mutation does not affect chemotaxis ability in the WT background but improves chemotaxis of nAβ animals (Fig. 4C). Imaging multi-neuron activity in these animals, we found that the *nmr-1* null mutation had no effect on positive neuron correlativity but increased the degree of neuron anti-correlativity in nAβ animals at adult day 6 (Fig. 4D, Table 4). In the WT background, *nmr-1* decreased positive correlativity and increased negative correlativity (Supp fig. 4b, Table 4) consistent with a loss of excitatory signaling.

## Discussion

### A unique loss of neuron connectivity in nAβ-mediated dysfunction

Our comprehensive neuronal imaging in aged *C. elegans* reveals a unique profile of functional decline resulting from amyloid beta (Aβ_1-42_) toxicity within a complete, intact nervous system. As illustrated in Figure 1, animals with pan-neuronal Aβ_1-42_expression exhibited a premature decline in system-wide activity dynamics, displaying highly stochastic and disorganized neuronal activity patterns in early adulthood, contrasting with the well-order neuronal dynamics of young wild-type animals. These discrepancies in network dynamics were reflected in behavioral deficits in mechanosensory as well as thermotactic and chemotactic responses of nAβ expressing animals. Within this context, we found that nAβ expressing animals displayed a loss of anti-correlated neuron connectivity similar to that of normal aging. Moreover, nAβ-expressing worms also exhibit a progressive loss of positive neuron connectivity as measured by a decline in the number of highly correlated neuron pairs. This is in contrast with normal aging in wild-type animals in which positive neuron correlations are maintained into old age (Wirak et al 2022). These findings were substantiated by the fact that nAβ expression also resulted in increased resistance to Aldicarb, suggesting decreased levels of ACh signaling in those animals. In mammals, Aβ toxicity has been shown to disrupt excitatory/inhibitory balance of the nervous system resulting in hyperexcitability within particular regions of the brain (33). A number of mammalian studies have also pointed to the vulnerability of ACh signaling in AD (34,35). Our findings here further suggest that a primary functional effect of Aβ toxicity on the nervous system is a generalized loss of excitatory connectivity that is unique to nAβ and distinct from the functional decline of normal aging.

### The role of one carbon metabolism (OCM) pathways in Aβ associated neuronal decline and its mitigation

Prior studies have reported metabolic stress is an early and primary events which affects AD pathology in transgenic *C. elegans* expressing pan neuronal human Aβ, with specific focus on energy metabolism (13). As evident from Figure 2, our findings point to the dysregulation of metabolic pathways in Aβ associated decline that are critical to synaptic function. nAβ expression resulted in downregulation of OCM-related genes *metr-1* and *sams-*1 that are involved in choline metabolism and could affect acetylcholine (ACh) availability. Conversely, supplementation with OCM metabolites (choline, methionine, and cysteine) in these animals helped rescue *metr-1* and *sams-*1 expression as well as the levels of ACh signaling (as assayed via aldicarb sensitivity). Analysis of the downstream transsulfuration pathway genes, that have also been clinically associated with neurodegenerative diseases (22,23) revealed possible disturbances into broader metabolic signaling cascades (46,47). At a functional level OCM supplements improved chemotactic behavior of aged nAβ expressing animals and cysteine in particular improved temporal nervous system dynamics. However, we did not measure recovery of Aβ mediated loss of positive neuron correlativity. Instead, OCM supplements improved anti-correlated neuron connectivity selectively in nAβ expressing animals while wild-type animals were largely un-affected.

Our investigation of the serine synthesis pathway (SSP) generated further evidence for the role of OCM in nAβ mediated functional decline. We manipulated the SSP either by blocking the enzymatic activity of the key SSP enzyme PHGDH *via* a PHGDH inhibitor NCT502 or by L-serine supplementation. As the SSP links glycolysis to OCM, these treatments should in turn modulate downstream choline metabolism by OCM. Many of our results support this hypothesis; blocking the SSP exacerbated both the loss of cholinergic signaling in nAβ animals (as measured by aldicarb resistance) as well as the number of highly correlated neuron pairs (measured via neuronal imaging). Likewise, L-serine supplementation enhanced cholinergic signaling in nAβ animals. Interestingly both treatments improved the degree of anti-correlated neuron signaling in old nAβ animals. At the molecular level, SSP manipulation helped to restore OCM gene expression in nAβ animals, suggesting the importance of metabolic crosstalk between glycolytic-serine flux and OCM pathways in AD. Paradoxically, any perturbation in SSP appeared to reduce ACh signaling in WT animals, increasing resistance to aldicarb and reducing positive neuronal correlativity. Such contradictory findings have, also, been reported in mammalian studies that differ as to the benefits of serine metabolism in AD (26,38). In an earlier study we reported similar contradictory effects of perturbation in the SSP on axonal regeneration in *C. elegans* (39). Clearly, while these metabolic pathways have important implications into Aβ associated neuronal decline and toxicity, such interventions also have context-dependent effects, necessitating careful evaluation before translation to therapeutic strategies.

Taken as a whole, these findings suggest that metabolic interventions of OCM and related pathways modulate aspects of neuronal decline but do not fully reverse Aβ-specific effects on neuronal connectivity. Indeed, the most consistent benefit appears to be an improvement in anti-correlated neuronal signaling that is lost during normal aging. These results speak to the importance and complexity of studying AD and Aβ pathology in the context of aging. These findings further suggest that potential therapeutics might mitigate the effects of the disease by improving the progression of normal aging. It is important to note that the bulk ensemble neuron activity measurements performed here may not reflect changes in critical subsets of neurons, such as the command interneurons that play an important role in system dynamics and behavior. In addition, it is possible that continued maintenance of the animals at 23°C throughout adulthood resulted in excessive overexpression of Aβ such that beneficial effects became imperceptible. For example, we measured no improvement in neuronal function with vitamin B12 supplementation despite previous studies linking it to neurodegenerative diseases (25,40). Regardless, our findings link OCM dysregulation to Aβ pathology at the metabolic, molecular and neurological levels, consistent with human evidence associating homocysteine and S-adenosylmethionine imbalances with cognitive decline in AD (22,41–43).

### Preservation of aging phenotypes in the context of Aβ-associated decline

We were able to mitigate the behavioral deficiency of Aβ expressing animals through direct manipulation of key synapse types within the *C. elegans* nervous system (Figure 4). We found that both *unc-2* gain-of-function mutation, which has been shown to lead to increased cholinergic signaling (29), and *nmr-1* null mutation, which reduces NMDAR activity implemented in Aβ-induced degeneration (44,45), effectively rescue the chemotaxis behavioral defect in nAβ expressing animals. However, rather than directly counteracting the specific effects of nAβ by preserving over-all positive neuron connectivity, these mutations helped to maintain anti-correlated connectivity of nAβ animals. Our inability to detect any benefit in the positive connectivity of the system could be due to the persistent ectopic nAβ expression that might mask relatively subtle effects. However, as with the metabolic manipulations, a major effect of these interventions appears to be the improvement of generalized aging phenotypes rather than specifically addressing the unique functional defects observed in nAβ expressing animals.

## Conclusion

*C. elegans* offers unparalleled access for *in vivo* imaging of neuronal activity and signaling across an entire nervous system with age. Applied in a nAβ expressing strain these capabilities allow for comprehensive measurement of Aβ induced insults on neuronal function with cellular resolution that have a direct impact on our understanding of the early stages of AD pathology. We find that Aβ toxicity results in pre-mature breakdown in system-wide neuron dynamics as well as a loss of positive, excitatory, neuron connectivity that is distinct from the effects of normal aging. Manipulation of metabolic pathways related to choline metabolism as well as key synapse types helped recovered behavioral phenotypes as well as some of the neurological effects of Aβ toxicity. Interestingly, these interventions also appeared to mitigate the loss of anti-correlated, inhibitory neuron signaling that is a function of normal, healthy aging, suggesting that the beneficial effects may stem in part from a protection against the processes of normal aging rather than directly mitigating the effects of Aβ. Our results, therefore, demonstrate both the unique aspects of Aβ toxicity on neuronal dynamics but also how they are related to, and dependent on, functional decline in normal aging.

## Methods Details

### KEY RESOURCES TABLE

| REAGENT | RESOURCE SOURCE | IDENTIFIER |
| --- | --- | --- |
| **Bacterial Strains** |  |  |
| E. coli OP50 | Caenorhabditis Genetics Centre | OP50 |
| **Chemicals and Pharmacological Reagents** |  |  |
| L-methionine | Fisher Scientific | AC166160025 |
| L-Cysteine | Sigma millipore | 168149-100G |
| Choline chloride | SANTA CRUZ BioTech | sc-207430B |
| L-Serine | Sigma | S-4500 |
| PHGDH Inhibitor (NCT502) | ChemScene LLC | CS-0064503 |
| Serotonin hydrochloride (5-TH) | SANTACRUZ Biotech | sc-201146A |
| β-mercaptoethanol | Fisher Scientific | AC125470100 |
| Trizol | Thermo Fisher Scientific | 15596018 |
| SYBR Green Master Mix | Applied Biosystems | A25741 |
| Sodium azide | Mallinckrodt | UN1687 |
| DMSO (Dimethyl Sulfoxide) | ThermoFisher Scientific | 85190, |
| Molecular Grade Water | invitrogen | 10977-015 500mL |
| Polyethylene glycol diacrylate | Advance BioMatrix | GS700 |
| Irgacure | Sigma-Aldrich | 410896 |
| Tetramisole | Sigma-Aldrich | L9756 |
| Cyanocobalamin (B12) | MilliPORE SiGMA | 47869 |
| Thiamine Hydrochloride (B1) | MilliPORE SiGMA | 47858 |
| Aldicarb | MilliPore Sigma | 33386-100MG |
| 5-Fluoro-2'-deoxyuridine | Santa Cruz | sc-202425B |
| Chloroform | Millipore Sigma | C2432-25ML |
| Thioflavine S | MilliPore Sigma | T1892-25G |
| Diacetyl | Sigma | S7584528237 |
| DMSO | MilliPore Sigma | D8418-50ML |
| DNAse-I | NEB | M03035 |
| Proteinase K | QIAGEN | 178010823 |

### REAGENTS AND RESOURCES

Further information and requests for resources, data and reagents should be directed to and will be fulfilled by the Lead Contact, Christopher V. Gabel.

### *C. elegans* strains maintenance

Hermaphrodite *C. elegans* were maintained on nematode growth medium (NGM)-agarose plates seeded Escherichia coli (strain OP50) as food source. Strains containing pan-neuronal expression of Human Amyloid Beta (Aβ_1-42_), CL2355 (smg-1(cc546) dvIs50 I.) (dvIs50 [pCL45 (snb-1::Abeta 1-42::3’ UTR(long) + mtl-2::GFP] I.) (14) were crossed with control imaging strains (QW1217;OH15263, see below). Aβ_1-42_ plaques were imaged using a strain with Human Aβ worms strain expression in muscular cells CL2006 (dvIs2 [pCL12(unc-54/human Abeta peptide 1-42 minigene) + rol-6(su1006)]). Other strains used in the study such as *unc-2*(gof), *nmr-1* etc. were also crossed in control imaging strain and nAβ expressing strain. Age synchronization was accomplished *via* bleaching, in which gravid adults were bleached on freshly prepared NGM plates. Experiments were performed on age matched day1, day3, and day6 of nematode adulthood. To maintain uniformity across the experiments all the worms were first bleached and maintained at 16°C till they reached L4/young adult (YA) stage, then they were transferred to higher temperature (either 20°C or 23°C) followed by behavior and imaging experiments on different days and conditions (Supp fig. 1a). Adult worms were regularly transferred to fresh plates, as needed, to prevent starvation and to separate the aging adults from their progeny, except when worms grow on plates containing 50 μM 5-fluoro-2′-deoxy uridine specifically for quantitative qRT-PCR and muscular Aβ fluorescence imaging experiments.

### Behavior Assays

All *C. elegans* behavior was assessed in a food-free environment. Touch response is performed for Day1, Day3 and Day6 to assess the temperature and age by which nAβ worms start exhibiting significant changes in behavior. For touch response, control worms (QW1217;OH15263) and nAβ (CL2355; QW1217;OH15263) worms were washed 3X in 1X S-Basal solution (100 mM NaCl, 50 mM KPO4 buffer, 5 μg/ml cholesterol) to remove bacteria before transferring to food-free NGM-agarose 35 mm plates, at room temperature, one worm per plate. Animals were then left undisturbed for 10 min on the food free plates before assaying. For anterior touch responsiveness individual animals, that were moving forward or not moving, were gently stroked with an eyelash across the anterior portion of their bodies and the response recorded. Animals were scored as responsive to a given stimulus, if a reversal movement was initiated. Each worm was subjected to touch response a total of 5 times, with a recovery period of at least 5 min between stimuli. A total of at least 20 worms were subjected to touch response assays and this was repeated 3 times, constituting a minimum 300 touch responses events across 60 animals per condition. For thermotaxis and chemotaxis assays, adult day6 worms were washed in S-basal 3X and then transferred to a food free NGM plate at 20°C for one hour to acclimatize before assaying.

For thermotaxis, ∼30-40 worms were placed in the center of a 90 mm plate on which a temperature gradient (20°C-23°C) was maintained. After one hour distribution of the worms was recorded and thermotaxis index was calculated according to the formula

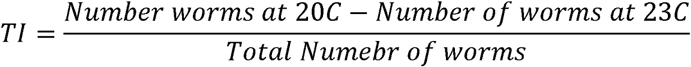

For the chemotaxis assay, 20-25 adult day 6 worms starved for 1-2 hours, were placed in the center of a 60 mm plate at 20°C. The plate was divided into quadrants, with diagonally opposite quadrants containing 2µl of the attractant 2% diacetyl (Sigma, cat# S7584528237, CAS:431-03-8) or 2µl 2% ethanol as control. Quadrants also contained 5mM sodium azide (Mallinckrodt, Cat# UN1687) to immobilize animals. After one hour distribution of the worms was scored by quadrant and the chemotaxis index was calculated according to the formula

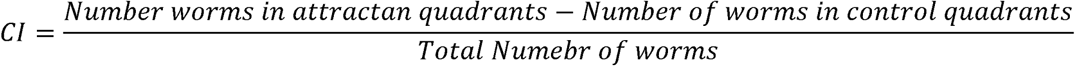

### Functional Multi Neuron Imaging

Light sheet multi neuronal functional imaging experiments were performed using the transgenic strains control, QW1217;OH15263 (lite-1[ce314], zfis146[nmr-1:: NLSwCherry::SL2::GCaMP6s, lim-4(− 3328–2174)::NLSwCherry::SL2::GCaMP6s, lgc-55 (−120–773)::NLSwCherry::SL2::GCaMP6s, npr-9:: NLSwCherry::SL2::GCaMP6s]. Which pan-neuronally expresses nuclear-localized tagRFP and cytoplasmic GCaMP6s and lite-1 mutant (QW1217) crossed with NeuroPal strain (OH15263) which contains four short pan-neuronal promoters fused together (unc-11::rgef-1::ehs-1::ric-19). NeuroPAL (Neuronal Polychromatic Atlas of Landmarks) transgene is used to resolve unique neural identities in whole-brain ganglion images. However, neuron identification proved prohibitively difficult in older and Aβ expressing animals and was not utilized in this study. Neuronal Aβ (nAβ) worm strain CL2355 (dvIs50 [pCL45 (snb-1::Abeta 1-42::3’ UTR(long) + mtl-2::GFP] I) was crossed with control imaging strain (QW1217;OH15263) to generate imaging strain for pan neuronal Aβ worms nAβ;QW;OH (CL2355; QW1217;OH15263). For all the imaging experiments age synchronized worms were cultured at 16°C till L4/young adult stage. At L4/YA stage control or nAβ worms were transferred to 23°C and initially worms were imaged at day1, day3 and day6. In case of drugs and metabolites supplementation, worms were transferred to plates containing drugs/metabolite at L4/YA stage and directly transferred to 23°C till the imaging experiments. For pan-neuronal functional imaging, sample preparation was performed as previously described (Awal et al., 2020). In brief, age synchronized worms are washed with S-basal (100 mM NaCl, 50 mM KPO4 buffer, 5 μg/ml cholesterol). Animals are then paralyzed in 20 μL of 5 mM tetramisole. The immobilized 5-10 worms were then encapsulated in 10 μL of a transparent and permeable polyethylene hydrogel on a silanized glass coverslip (prepared as described; Burnett et al., 2018). Following hydrogel encapsulation, worms were placed in a Petri dish (90 mm) and immersed in S-Basal solution (100 mM NaCl, 50 mM KPO4 buffer, 5 μg/ml cholesterol) containing 5 mM tetramisole. Imaging was performed using a dual-inverted selective plane illumination fluorescence microscope (Applied Scientific Instrumentation, USA) with water immersion 0.8 NA 40. objectives (Nikon USA, Melville, NY). Each animal was imaged for 10 min at a rate of two volumes/second (voxel size 0.1625. 0.1625. 1μm), capturing both nuclear-RFP and cytoplasmic-GCaMP6s fluorescence. Custom Python scripts were employed during postprocessing to track 120 RFP-labeled nuclei per animal in three dimensions and extract GCaMP6s signals from the surrounding somas as described in Wirak et. al (11).

### Functional Multi Neuron Imaging Data analysis

For multi-neuron activity imaging each condition consists of an average of 12-20 worms, with activity traces measured from 120 neurons per animal via custom image analysis algorithms described in Awal et al., 2020. Normalized fluorescence, ΔF/F_0_, of all measured neurons was plotted in the activity heatmap for each condition as shown in Figure 1 D-F left panels. For each neuron, F_o_ was calculated as the mean value of the lowest 1% of measurements made for that neuron. Neuron activity heatmaps were scaled to the dynamic range of values in that trial. Principle component analysis (PCA) and neuronal correlation analysis were performed as in Wirak et al 2022. In brief, the first three principal components were plotted on a 3D graph to generate a trajectory over time, Figure 1 D-F center panels. Smoothness of these trajectories was measured by calculating the instantaneous change in direction at each time point, *i.e.* the absolute angular difference between the tangential direction over the past three seconds and that of the future three seconds. Probability histograms for individual animals (trials) or across all trials of a specific condition were generated by pooling the measurements of angular change from all timepoints within that/those trial(s), Figure 1 D-F right, bottom panels. Neuron correlativity was calculated as the Pearson correlation between the differentiated signals from each possible neuron pair among the 40 most dynamically active neurons (i.e. those having the greatest standard deviation in neuronal signal) of the 120-neurons measured in each animal, Figure 1 D-F right top, panels.

### Drug and metabolite treatments in *C. elegans*

For all pharmacological reagents and metabolite treatments, the compound was dissolved and supplemented in NGM agar before plate preparation and stored at 4°C. For all supplementation experiments animals were cultured on treated plates from L4/YA stage till the time of assay. Choline 30 mM (Sigma, Cat#: C7017-5G, CAS: 67-48-1) (46), L-Methionine 10 mM (Fisher Scientific, Cat#: AC166160025), 5 mM L-Serine (Sigma, cat# S-4500) (Liu et al., 2019), L-Cysteine 10 mM (Sigma, Cat#:168149-25G, CAS:52-90-4), were dissolved in molecular grade water (Fisher Scientific, Cat#: R91450001G, CAS: 7732-18-5) at required stock concentrations (39). The phgdh-1 (C31C9.2) inhibitor N-(4,6-dimethylpyridin-2-yl)–4-[5-(trifluoromethyl)yridine-2-yl] piperazine-1-carbothioamid (NCT502) (MedChemExpress, Cat#: HY-117240) was initially dissolved in DMSO and diluted in ddH2O to use at a concentration of 25 μM in NGM plates as reported earlier (39). Aldicarb (Sigma, Cat#:33386-100MG, CAS:116-06-3), 100 mM aldicarb stock solution was prepared by dissolving 100 mg of aldicarb in 5.25 ml of 70% ethanol, store at -20 °C, and final concentration of 1 mM was used in NGM plate for the assay. All the drugs, chemicals, and metabolites used are listed in the KEY RESOURCES TABLE at the beginning of method details.

### Aldicarb Paralysis Assay

The Aldicarb sensitivity assay was performed to assess neuromuscular function as proxy for cholinergic neuronal signaling in different *C. elegans* strains (47) Aldicarb (O-(Methylcarbamoyl)-2-methyl-2-(methylthio)-propionaldehyd-oxime) (Sigma-Aldrich, SKU# 33387-100MG) stock solution (100 mM) was prepared in 70% ethanol and diluted to a final concentration of 1 mM in molten NGM before pouring into assay plates. Plates were dried for 24 hours at room temperature before use. For the assay, approximately 20-30 adult day 6 worms were transferred to aldicarb-containing plates at room temperature and observed for paralysis at 20-minute intervals for up to 2-3 hours. Paralysis was defined as immobility and the inability to respond to gentle poking with a platinum wire. Three independent trials were conducted for each condition, and results were expressed as the percentage of paralyzed worms over time.

### Fluorescence staining of muscular Aβ deposits

Individual control (QW1217;OH15263) and transgenic *C. elegans* expressing human amyloid beta peptide in muscles (CL2006) were fixed in 4% paraformaldehyde/PBS, pH 7.4, for 24 h at 4°C, and permeabilized in 5% fresh β-mercaptoethanol, 1% Triton X-100, 125 mM Tris, pH 7.4, in a 37°C incubator for 24 h. The nematodes were stained for Aβ plaques with 0.125% thioflavin S (Sigma) in 50% ethanol for 2 minutes, destained by 5X washing with S-basal buffer, and mounted on 2% agarose pad slides for microscopy (14). Fluorescence images were acquired at consistent exposure parameters using a 40X objective of the BZ X800 - KEYENCE microscope. The number of thioflavin S-reactive deposits in the anterior area of the pharyngeal bulb in individual animals were scored.

### qRT PCR

To evaluate the expression levels of candidate genes under different experimental conditions and strains, we employed quantitative real time polymerase chain reaction (qRT-PCR). Animals were grown on plates containing 50 μM 5-fluoro-2′-deoxy uridine. Day 6 adult worms were lysed in 0.5% SDS, 5% beta-ME, 10 mM EDTA, 10 mM Tris-HCl pH 7.4, 0.5 mg/ml Proteinase K, and RNA was purified with Tri-Reagent (Sigma). 2–3 μg RNA was subjected to DNAse I treatment (NEB M03035), followed by cDNA conversion using High-Capacity cDNA Reverse Transcription Kit (Thermo Fisher Scientific cat#4368814). qRT-PCR was performed in biological triplicate with two technical duplicates for each condition using Applied Biosciences Real-Time PCR Machine and Fast SYBR Green Master Mix (Thermo Fisher, 4385617). Relative transcript abundance was determined by using the DDCt method and normalized to *act-1* mRNA expression levels as a control. All the qRT-PCR Primers used in this study are listed in Table5.

### Statistical analysis

For all behavior assays and qRT-PCR data unpaired T-test or One-way ANOVA analyses with Bonferroni’s Multiple Comparison Test were performed to assess the statistical significance between the groups. Nonparametric, Mann Whitney test was applied to evaluate the statistical differences in positively, and negatively, correlated neuron pairs across conditions. The value of p ≤0.05 is considered statistically significant. MATLAB, GraphPad, and Microsoft excel were used for analysis and figure generation.

## Additional data information

Table1_Data_Figure 1

Table2_Data_Figure 2

Table3_Data_Figure 3

Table4_Data_Figure 4

Table5_RT-PCR_Primers

## Author Contribution

DKY and CVG study conceptualization with CWC, DKY data acquisition, DKY, CWC and CVG formal analysis, DKY, CWC and CVG Methodology, DKY designed and performed experiments, and data analysis with the help of CWC and CVG, writing of original draft. CWC and CVG contributed to the final draft with input from all the authors.

## Supplemental Figure

**Supp fig. 1.**
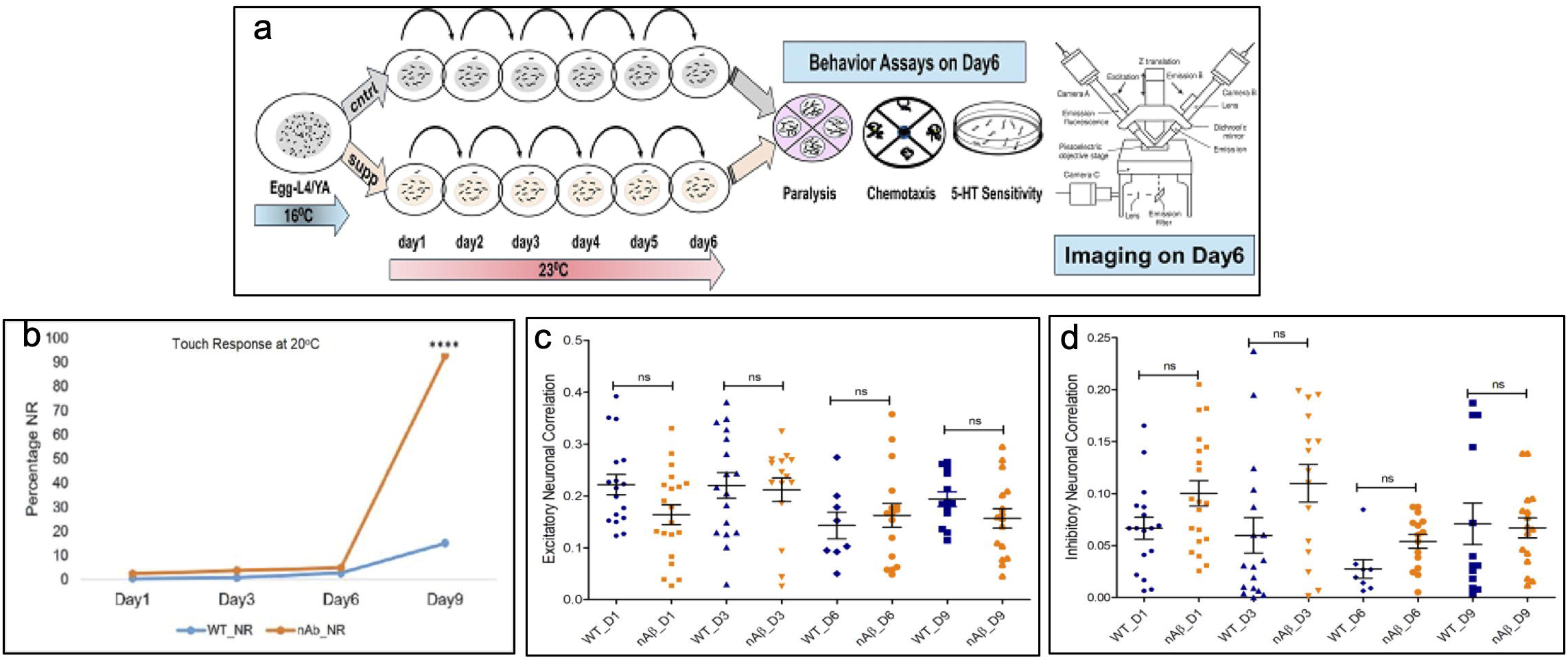
(a) Schematic representation of experimental paradigm demonstrating the worms cultures/treatments for behavior analysis and multi-neuron imaging. (b) Anterior touch response (percent not responsive) WT (blue) and nAβ (orange) for worms reared at 20°C from young adult/day 1 till day 9. (c) Quantitative comparison of excitatory signaling in WT (blue) and nAβ (orange) worms at 20°C on day1, day3, day6 and day9. (d) Quantitative comparison of inhibitory signaling in WT and nAβ worms at 20°C on day1, day3, day6 and day9.

**Supp fig. 2.**
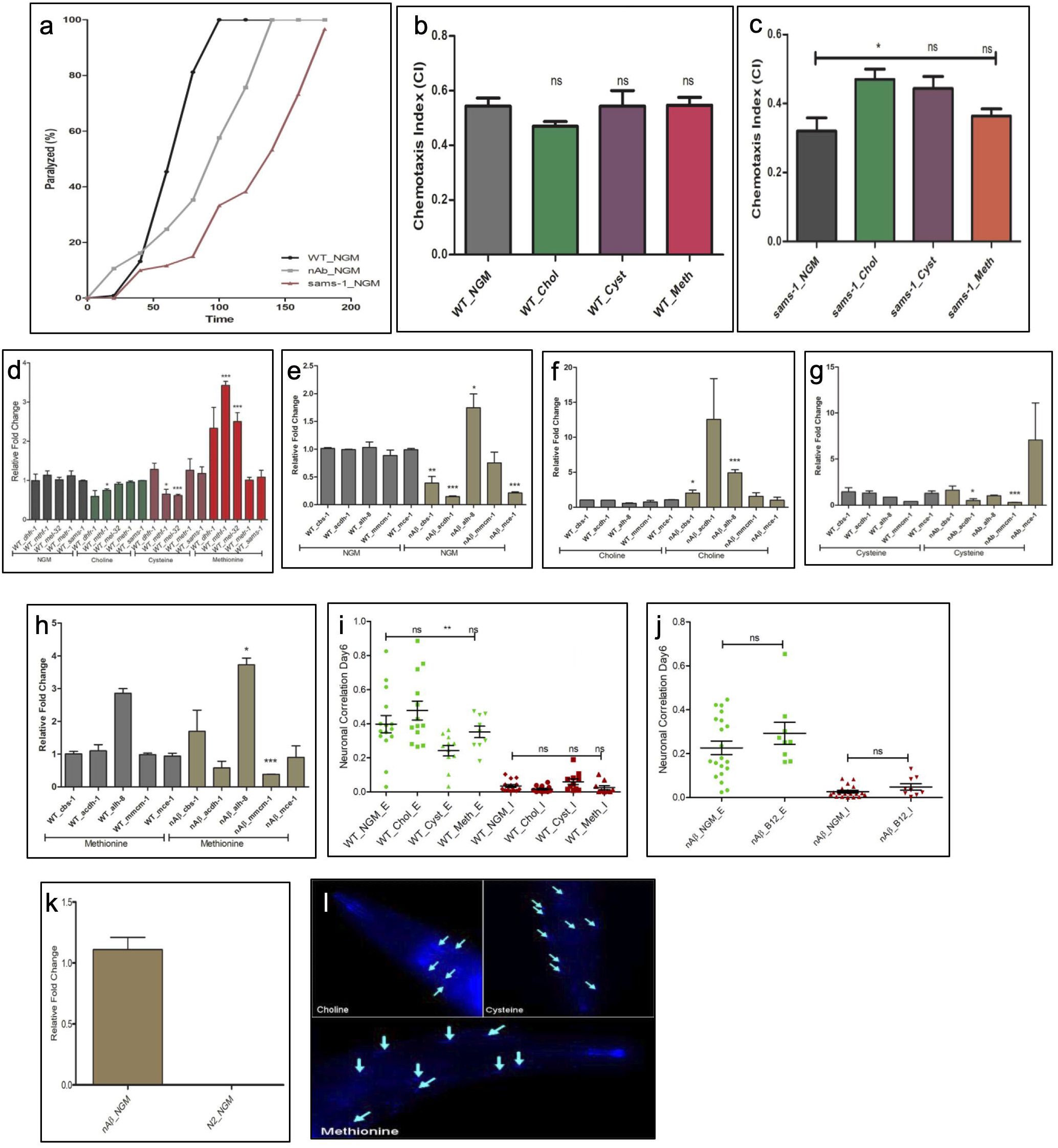
(a) Acetyl choline signaling assessed by aldicarb assay in WT, nAβ and *sams-1* mutant worms. (b) Effect of OCM metabolite (choline, cysteine and methionine) supplementation on chemotaxis behavior assay in WT worms. (c) Effect of OCM metabolite (choline, cysteine and methionine) supplementation on chemotaxis behavior assay in *sams-1* worms. (d) Effect of Choline, Cysteine and Methionine supplementation on the expression level of selected genes critical for OCM pathway in WT worms. (e) Expression level of selected genes in transulfuration and downstream pathway in WT and nAβ worms. Effect of (f) Choline, (fg Cysteine and (h) Methionine supplementation on the expression level of selected genes in transulfuration and downstream pathway in control and nAβ worms. (i) Quantitative comparison of excitatory (green) and inhibitory (red) signaling in WT worms supplemented with OCM metabolites. (j) Quantitative comparison of excitatory (green) and inhibitory (red) signaling in nAβ worms supplemented with Vit. B12. (k) Human amyloid beta trans gene (Aβ_1–42_) expression level in nAβ worms and WT worms. (l) Representative images showing ThS stained *C. elegans* that express the Aβ_1–42_ in muscle cell (CL2006) supplemented with OCM metabolites in the pharyngeal region. Measurements were done at adult day 6.

**Supp fig 3.**
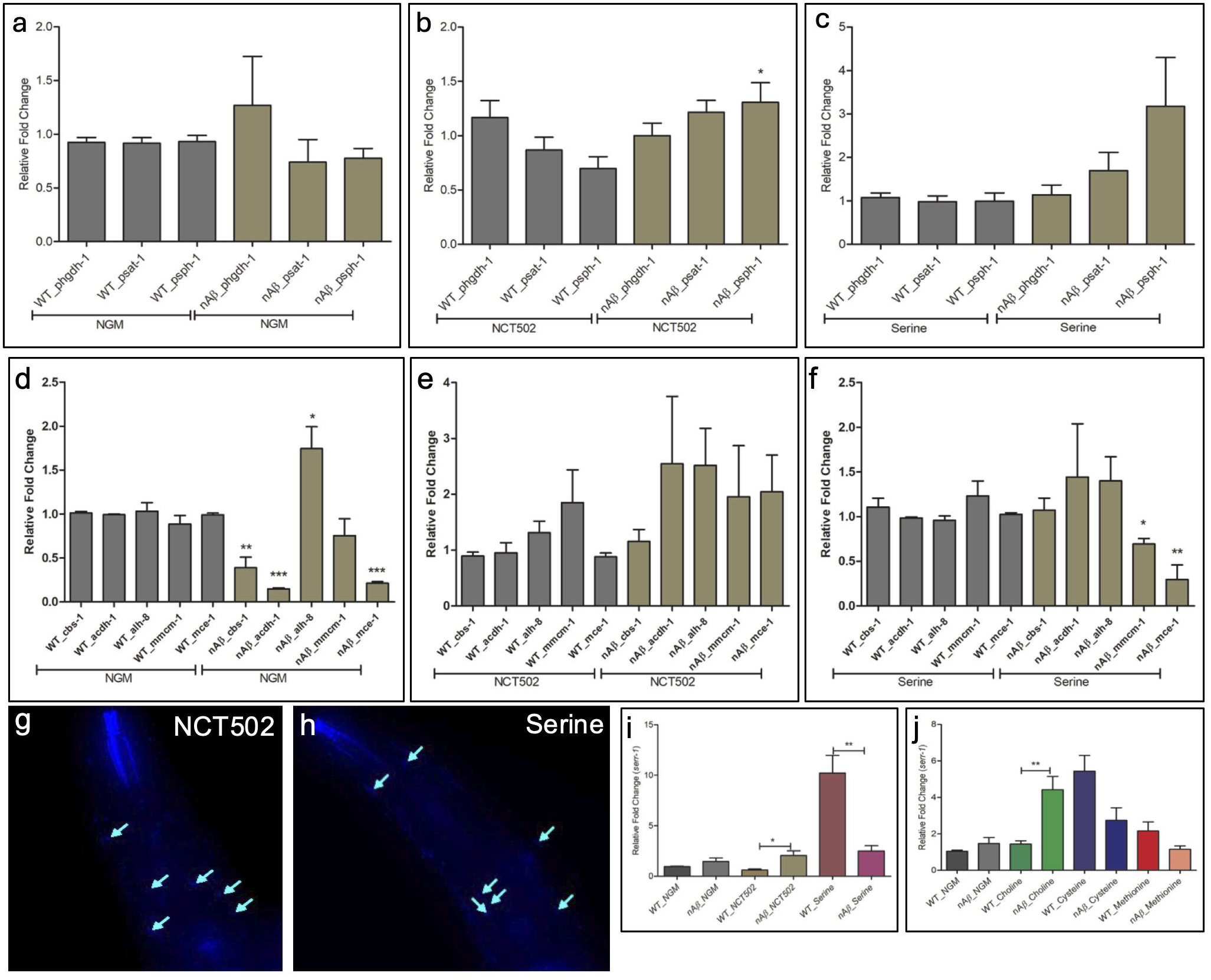
(a) Expression level of phdh-1, psat-1 and psph-1 genes in WT and nAβ worms. Expression level of phdh-1, psat-1 and psph-1 genes in WT and nAβ worms upon (b) NCT502 and (c) serine supplementation. (d) Expression level of *cbs-1, acdh-1, alh-8, mmcm-1* and *mce-1* genes in WT and nAβ worms. Expression level of *cbs-1, acdh-1, alh-8, mmcm-1* and *mce-1* genes in WT and nAβ worms upon (e) NCT502 supplementation and (f) upon serine supplementation. Images showing ThS stained *C. elegans* that express the Aβ_1–42_ in muscle cell (CL2006) supplemented with (g) NCT502 and (h) serine in the pharyngeal region. (i) Expression level of *serr-1* gene in WT and nAβ worms upon NCT502 and Serine supplementation. (j) Expression level of *serr-1* gene in WT and nAβ worms upon OCM metabolites supplementation. All animals were tested at adult day 6.

**Supp fig 4.**
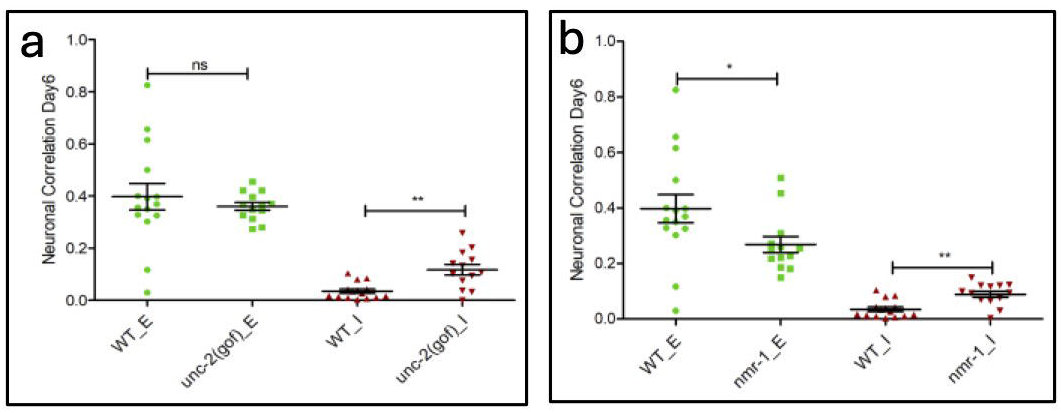
(a) Quantitative comparison of excitatory (green) and inhibitory (red) neuronal signaling in WT and unc-2(gof) worms at adult day6. (b) Quantitative comparison of excitatory (green) and inhibitory (red) signaling in WT and *nmr-1* worms at adult day6. Quantitative comparison of excitatory (green) and inhibitory (red) signaling in WT and spr-4 worms at adult (c) day1, (d) day3, and (e) day6 old. (j) Quantitative comparison of excitatory (green) and inhibitory (red) signaling in nAβ and nAβ supplemented with X5050 worms at adult (f) day1, (g) day3 and (h) day6 old.

## Supplemental Tables

**Table1_Data_Figure 1 and Supp fig. 1**.

**Table2_Data_Figure 2 and Supp fig. 3**.

**Table3_Data_Figure 3 and Supp fig. 3**.

**Table4_Data_Figure 4 and Supp fig. 4**.

**Table5_RT-PCR_Primers**

## References

1. Holtzman DM, Morris JC, Goate AM. Alzheimer’s disease: The challenge of the second century. Vol. 3, Science Translational Medicine. 2011.

2. Shankar GM, Walsh DM. Alzheimer’s disease: Synaptic dysfunction and Aβ. Mol Neurodegener. 2009;4(1).

3. Frere S, Slutsky I. Alzheimer’s Disease: From Firing Instability to Homeostasis Network Collapse. Vol. 97, Neuron. 2018.

4. Liu Y, Tan Y, Zhang Z, Yi M, Zhu L, Peng W. The interaction between ageing and Alzheimer’s disease: insights from the hallmarks of ageing. Vol. 13, Translational Neurodegeneration. 2024.

5. Pena F, Gutierrez-Lerma A, Quiroz-Baez R, Arias C. The Role of beta-Amyloid Protein in Synaptic Function: Implications for Alzheimers Disease Therapy. Curr Neuropharmacol. 2006;4(2).

6. Mucke L, Selkoe DJ. Neurotoxicity of amyloid β-protein: Synaptic and network dysfunction. Cold Spring Harb Perspect Med. 2012;2(7).

7. Guo T, Zhang D, Zeng Y, Huang TY, Xu H, Zhao Y. Molecular and cellular mechanisms underlying the pathogenesis of Alzheimer’s disease. Vol. 15, Molecular Neurodegeneration. 2020.

8. Coupé P, Manjón JV, Lanuza E, Catheline G. Lifespan Changes of the Human Brain In Alzheimer’s Disease. Sci Rep. 2019;9(1).

9. Roussos A, Kitopoulou K, Borbolis F, Palikaras K. Caenorhabditis elegans as a Model System to Study Human Neurodegenerative Disorders. Vol. 13, Biomolecules. 2023.

10. Awal MR, Wirak GS, Gabel C V., Connor CW. Collapse of Global Neuronal States in Caenorhabditis elegans under Isoflurane Anesthesia. Anesthesiology. 2020 Jul 1;133(1):133–44.

11. Wirak GS, Florman J, Alkema MJ, Connor CW, Gabel C V. Age-associated changes to neuronal dynamics involve a disruption of excitatory/inhibitory balance in C. elegans. Elife. 2022;11.

12. Maestú F, de Haan W, Busche MA, DeFelipe J. Neuronal excitation/inhibition imbalance: core element of a translational perspective on Alzheimer pathophysiology. Vol. 69, Ageing Research Reviews. 2021.

13. Teo E, Ravi S, Barardo D, Kim HS, Fong S, Gassiot AC, et al. Metabolic stress is a primary pathogenic event in transgenic Caenorhabditis elegans expressing pan-neuronal human amyloid beta. Elife. 2019;8.

14. Wu Y, Wu Z, Butko P, Christen Y, Lambert MP, Klein WL, et al. Amyloid-β-induced pathological behaviors are suppressed by Ginkgo biloba extract EGB 761 and ginkgolides in transgenic Caenorhabditis elegans. Journal of Neuroscience. 2006;26(50).

15. Kato S, Kaplan HS, Schrödel T, Skora S, Lindsay TH, Yemini E, et al. Global Brain Dynamics Embed the Motor Command Sequence of Caenorhabditis elegans. Cell. 2015;163(3).

16. Rand JB. Acetylcholine (January 30, 2007). In: WormBook, ed The C elegans Research Community, WormBook, doi/101895/wormbook11311, http://www.wormbook.org. 2007.

17. Mahoney TR, Luo S, Nonet ML. Analysis of synaptic transmission in Caenorhabditis elegans using an aldicarb-sensitivity assay. Nat Protoc. 2006;1(4).

18. Zeisel SH. Metabolic crosstalk between choline/1-carbon metabolism and energy homeostasis. In: Clinical Chemistry and Laboratory Medicine. 2013.

19. Borro M, Cavallaro RA, Gentile G, Nicolia V, Fuso A, Simmaco M, et al. One-carbon metabolism alteration affects brain proteome profile in a mouse model of Alzheimer’s disease. Journal of Alzheimer’s Disease. 2010;22(4).

20. Joshi SM, Jadavji NM. Deficiencies in one-carbon metabolism led to increased neurological disease risk and worse outcome: homocysteine is a marker of disease state. Vol. 11, Frontiers in Nutrition. 2024.

21. Lionaki E, Ploumi C, Tavernarakis N. One-Carbon Metabolism: Pulling the Strings behind Aging and Neurodegeneration. Vol. 11, Cells. 2022.

22. Figueroa-González G, Silva-Adaya D, Corona-Trejo A, Gonsebatt ME, Trejo-Solis C, Campos-Peña V, et al. Transsulfuration pathway: a targeting neuromodulator in Parkinson’s disease. Rev Neurosci. 2023;34(8).

23. Paul BD. Neuroprotective Roles of the Reverse Transsulfuration Pathway in Alzheimer’s Disease. Vol. 13, Frontiers in Aging Neuroscience. 2021.

24. Umekar M, Premchandani T, Tatode A, Qutub M, Raut N, Taksande J, et al. Vitamin B12 deficiency and cognitive impairment: A comprehensive review of neurological impact. Vol. 18, Brain Disorders. Elsevier B.V.; 2025.

25. Luthra NS, Marcus AH, Hills NK, Christine CW. Vitamin B12 measurements across neurodegenerative disorders. J Clin Mov Disord. 2020;7(1).

26. Bonvento G, Oliet SHR, Panatier A. Glycolysis-derived L-serine levels versus PHGDH expression in Alzheimer’s disease. Vol. 34, Cell Metabolism. 2022.

27. Le Douce J, Maugard M, Veran J, Matos M, Jégo P, Vigneron PA, et al. Impairment of Glycolysis-Derived L-Serine Production in Astrocytes Contributes to Cognitive Deficits in Alzheimer’s Disease. Cell Metab. 2020;31(3).

28. Ni X, Inoue R, Wu Y, Yoshida T, Yaku K, Nakagawa T, et al. Regional contributions of D-serine to Alzheimer’s disease pathology in male App^NL–G–F/NL–G–F^ mice. Front Aging Neurosci. 2023;15.

29. Huang YC, Pirri JK, Rayes D, Gao S, Mulcahy B, Grant J, et al. Gain-of-function mutations in the UNC-2/CaV2α channel lead to excitation-dominant synaptic transmission in C. elegans. Elife. 2019;8.

30. Bukke VN, Archana M, Villani R, Romano AD, Wawrzyniak A, Balawender K, et al. The dual role of glutamatergic neurotransmission in Alzheimer’s disease: From pathophysiology to pharmacotherapy. Vol. 21, International Journal of Molecular Sciences. 2020.

31. Liu J, Chang L, Song Y, Li H, Wu Y. The role of NMDA receptors in Alzheimer’s disease. Vol. 13, Frontiers in Neuroscience. 2019.

32. Liu W, Li Y, Zhao T, Gong M, Wang X, Zhang Y, et al. The role of N-methyl-D-aspartate glutamate receptors in Alzheimer’s disease: From pathophysiology to therapeutic approaches. Vol. 231, Progress in Neurobiology. 2023.

33. Nelson SB, Valakh V. Excitatory/Inhibitory Balance and Circuit Homeostasis in Autism Spectrum Disorders. Vol. 87, Neuron. 2015.

34. Hampel H, Mesulam MM, Cuello AC, Khachaturian AS, Vergallo A, Farlow MR, et al. Revisiting the Cholinergic Hypothesis in Alzheimer’s Disease: Emerging Evidence from Translational and Clinical Research. Vol. 6, The journal of prevention of Alzheimer’s disease. 2019.

35. Chen ZR, Huang JB, Yang SL, Hong FF. Role of Cholinergic Signaling in Alzheimer’s Disease. Vol. 27, Molecules. 2022.

36. Ionescu-Tucker A, Cotman CW. Emerging roles of oxidative stress in brain aging and Alzheimer’s disease. Vol. 107, Neurobiology of Aging. 2021.

37. Obeid R, Herrmann W. Mechanisms of homocysteine neurotoxicity in neurodegenerative diseases with special reference to dementia. Vol. 580, FEBS Letters. 2006.

38. Chen X, Calandrelli R, Girardini J, Yan Z, Tan Z, Xu X, et al. PHGDH expression increases with progression of Alzheimer’s disease pathology and symptoms. Vol. 34, Cell Metabolism. 2022.

39. Yadav DK, Chang AC, Grooms NW, Chung SH, Gabel C V. O-GlcNAc signaling increases neuron regeneration through one-carbon metabolism in Caenorhabditis elegans. Elife. 2024 Feb 9;13.

40. Umekar M, Premchandani T, Tatode A, Qutub M, Raut N, Taksande J, et al. Vitamin B12 deficiency and cognitive impairment: A comprehensive review of neurological impact. Vol. 18, Brain Disorders. Elsevier B.V.; 2025.

41. Tawfik A, Elsherbiny NM, Zaidi Y, Rajpurohit P. Homocysteine and age-related central nervous system diseases: Role of inflammation. Vol. 22, International Journal of Molecular Sciences. 2021.

42. Luzzi S, Cherubini V, Falsetti L, Viticchi G, Silvestrini M, Toraldo A. Homocysteine, Cognitive Functions, and Degenerative Dementias: State of the Art. Vol. 10, Biomedicines. 2022.

43. Hooshmand B, Refsum H, Smith AD, Kalpouzos G, Mangialasche F, Von Arnim CAF, et al. Association of Methionine to Homocysteine Status with Brain Magnetic Resonance Imaging Measures and Risk of Dementia. JAMA Psychiatry. 2019;76(11).

44. Tackenberg C, Grinschgl S, Trutzel A, Santuccione AC, Frey MC, Konietzko U, et al. NMDA receptor subunit composition determines beta-amyloid-induced neurodegeneration and synaptic loss. Cell Death Dis. 2013;4(4).

45. Gu Z, Cheng J, Zhong P, Qin L, Liu W, Yan Z. Aβ selectively impairs mGluR7 modulation of NMDA signaling in basal forebrain cholinergic neurons: Implication in alzheimer’s disease. Journal of Neuroscience. 2014;34(41).

46. Smulan LJ, Ding W, Freinkman E, Gujja S, Edwards YJK, Walker AK. Cholesterol-Independent SREBP-1 Maturation Is Linked to ARF1 Inactivation. Cell Rep. 2016;16(1).

47. Oh K, Kim H. Aldicarb-induced Paralysis Assay to Determine Defects in Synaptic Transmission in Caenorhabditis elegans. Bio Protoc. 2017;7(14).

